# A non-invasive digital biomarker for the detection of rest disturbances in the SOD1G93A mouse model of ALS

**DOI:** 10.1101/2019.12.27.889246

**Authors:** E. Golini, M. Rigamonti, F. Iannello, C. De Rosa, F. Scavizzi, M. Raspa, S. Mandillo

## Abstract

Amyotrophic Lateral Sclerosis (ALS) is a devastating neurodegenerative disease that affects both central and peripheral nervous system, leading to the degeneration of motor neurons, which eventually results in muscle atrophy, paralysis and death. Sleep disturbances are common in patients with ALS, leading to even further deteriorated quality of life. Investigating methods to potentially assess sleep and rest disturbances in animal models of ALS is thus of crucial interest.

We used an automated home cage monitoring system (DVC^®^) to capture activity patterns that can potentially be associated with sleep and rest disturbances and thus to the progression of ALS in the SOD1G93A mouse model. DVC^®^ enables non-intrusive 24/7 long term animal activity monitoring, which we assessed together with body weight decline and neuromuscular function deterioration measured by grid hanging and grip strength tests in male and female mice from 7 until 24 weeks of age.

We show that as the ALS progresses over time in SOD1G93A mice, activity patterns during day time start becoming irregular, with frequent activity bouts that are neither observed in control mice nor in SOD1G93A at a younger age. The increasing irregularities of activity patterns during day time are quantitatively captured by designing a novel digital biomarker, referred to as Rest Disturbance Index (RDI). We show that RDI is a robust measure capable of detecting rest/sleep-related disturbances during the disease progression earlier than conventional methods, such as the grid hanging test. Moreover RDI highly correlates with grid hanging and body weight decline, especially in males.

The non-intrusive long-term continuous monitoring of animal activity enabled by DVC^®^ has been instrumental in discovering activity patterns potentially correlated with sleep and rest disturbances in the SOD1G93A mouse model of the ALS disease.

## Introduction

Amyotrophic Lateral Sclerosis (ALS) is a devastating neurodegenerative disease that affects both central and peripheral nervous system, and it is characterized by the degeneration of upper and lower motor neurons that will result in muscle atrophy, paralysis and death within 2-5 years after diagnosis [1]. Death usually comes from respiratory failure. Along with progressive voluntary skeletal muscle weakness and atrophy, symptoms include dysphagia, dysarthria, respiratory dysfunction and sleep disturbances. Sleep disruption is very common in ALS and it is often related to hypoventilation, hypoxia, hypercapnia, restless legs, immobilization, nocturnal cramps and pain [2, 3]. Sleep disturbances are also very common in other neurodegenerative diseases, as Alzheimer’s, Parkinson’s and have an enormous impact on the quality of life of patients [4-6]. Circadian and sleep dysfunctions are often premorbid and can serve as early diagnostic markers of neurodegeneration.

Some studies in animal models of neurodegenerative diseases tackle this important issue [7, 8] but few are specific for ALS models. It is therefore crucial to provide evidence of this early symptom and to propose a suitable tool to investigate circadian and sleep related disruption in a classical mouse model of ALS as the SOD1G93A transgenic strain [9]. The SOD1G93A mouse model we used expresses multiple copies of the human mutated form of the SOD1 gene (on a C57BL/6J background) and it recapitulates the progressive symptomatology of the disease starting around 14-16 weeks of age: muscle weakness, tremors, body weight loss, limb paralysis, respiratory failure and death around 160-170 days or 24-25 weeks of age and it also presents ALS characteristic neurobiological substrates (cortical and spinal cord motor neuron degeneration, gliosis, neuroinflammation) [10, 11].

To study activity and rest in the SOD1G93A mouse model during the pre-symptomatic and symptomatic stages of the disease we adopted a technology capable of monitoring animal activity directly in the home cage [12-14]. Such systems are generally designed with the aim of collecting animal activity data 24/7, without interfering with nor handling the animals. This has the potential to unveil animal behaviors occurring at any time during the day [13] and over extended observation periods ranging from days to months and years [14]. The application of 24/7 long term home cage animal monitoring is particularly promising in the field of neurodegenerative diseases, especially the ones that potentially manifest complex modifications of the activity behaviors as the disease progresses over time, and that can be potentially hard to capture when observing animals outside the home cage as in conventional testing procedures (e.g., open field, rotarod, grid hanging ect.). In this experiment we monitored animals for several months, using the home cage monitoring system, referred to as Tecniplast DVC^®^ [15], which enables automated and non-intrusive long term data collection.

The main goal of this paper is to explore and better understand disease progression over time while keeping animals in a familiar environment (i.e., the home cage) for most of their life. This may lead to unveil activity patterns that would be hardly measurable otherwise. Particularly, we developed a new digital biomarker to detect rest disturbances during day time by observing animals for extended periods. In addition to home cage monitoring, we also performed tests commonly used in assessing ALS-related symptoms, such as body weight loss and neuromuscular decline (grid hanging and grip strength tests). We demonstrated that the use of home cage monitoring allowed us to capture rest disturbances at early stages of the disease. At the symptomatic stage we also showed that peak of rest disturbances are in good correlation with other symptoms of the disease.

## Materials and Methods

### Subjects

Male B6.Cg-Tg(SOD1*G93A)1Gur/J mice carrying a high copy number of mutant human SOD1 allele [9] were purchased at the Jackson Laboratories (Bar Harbor, ME, USA, stock n. 004435) and a mouse colony was established in-house at the CNR-EMMA-Infrafrontier-IMPC facility (Monterotondo, Italy) by crossing hemizygous transgenic males with C57BL/6J females. Progeny was genotyped by standard PCR following the Jackson Lab protocol (www.jax.org). Litter- and sex-matched pups were raised group-housed in standard cages (Thoren, Hazleton, PA, USA) enriched with a transparent red polycarbonate igloo house (Datesand, Manchester, UK) and with wood shavings contained in single cellulose bags (Scobis Uno bags, Mucedola, Settimo Milanese, Italy). Food (Standard Diet 4RF21, Mucedola, Italy) and water were available *ad libitum*. Room temperature was 21±2°C, relative humidity was 50-60%, and mice were kept in a 12h light/dark cycle with lights on at 07:00 am until 07:00 pm. Animals were subjected to experimental protocols approved by the Local Animal Welfare Committee and the Veterinary Dept. of the Italian Ministry of Health (Aut. #914/2016-PR), and experiments were conducted according to the ethical and safety rules and guidelines for the use of animals in biomedical research provided by the relevant Italian laws and European Union’s directives (Italian Legislative Decree 26/2014 and 2010/63/EU). All adequate measures were taken to minimize animal pain or discomfort. Extra wet food was provided inside the cage as needed.

### Experimental design

Male and female wild-type (WT) and transgenic (SOD1G93A) littermate mice at the age of 7 weeks were transferred to the DVC^®^ rack (see below), housing two mice per cage of same sex and genotype and assigned to the following experimental groups: 1) Males, WT n=10 (5 cages); 2) Males, SOD1G93A (TG) n=10 (5 cages); 3) Females, WT n=16 (8 cages); 4) Females, SOD1G93A (TG) n=12 (6 cages). Experiments have been conducted over two years, testing two separate cohorts of mice (Cohort I N=18, Cohort II N= 30).

From the age of 7 weeks, all mice were weighed weekly and tested for neuromuscular function using the grid hanging (weekly) and grip strength (every two weeks only for the cohort II) tests. We present the grip strength results in the Supporting Information for reference since this test has been performed on cohort II only. Mice of the first cohort were sacrificed at 22 weeks of age based on failure on the grid hanging test. Mice of the second cohort were monitored until humane endpoint (body weight loss >20% or loss of righting reflex), around age 24 weeks, and thus sacrificed according to current laws and regulations. Cages were changed every two weeks for cohort I (due to mouse facility management priorities) and once a week for cohort II, we verified that this difference did not affect data outcome.

### Home cage activity monitoring: Digital Ventilated Cage (DVC^®^) system

All mice were housed in a Digital Ventilated Cage (DVC^®^) rack, which is equipped with a home cage monitoring system capable of automatically measuring animal activity 24/7 [15]. DVC^®^ rack is installed on a standard IVC rack (Tecniplast DGM500, Buguggiate, Italy) by adding sensing technologies externally to the cage, so that neither modifications nor intrusion occur in the home cage environment. Animal locomotion activity is monitored via a capacitance sensing technology by means of 12 contactless electrodes, uniformly distributed underneath the cage floor, which record animal movements based on their presence in each electrode surrounding. In this work, animal activity is captured similarly to the *activation density* metric defined in a previous study [15], and by choosing to aggregate measurements in bins of 1-minute (*raw activity* data). For simplicity, we refer to this metric as *activity* throughout the paper. Since this study lasted several months (compared to the 1-minute binning), we decided to condensate the 1-minute raw activity into 2 distinct measurements per week: *i*) average activity during night time (i.e., by averaging all 1-minute bins within night period in each week); *ii*) average activity during day time.

### Rest Disturbance Index (RDI)

In this paper, we introduce the use of a novel digital biomarker, referred to as Rest Disturbance Index (RDI), which has been developed to capture irregular animal activity patterns during resting periods (day time). To quantitatively capture these patterns, we designed RDI based on the sample entropy [16] as the core metric, in which we set parameters m=2, r=0.2 and N=720 (as per notation in [16]). We computed the sample entropy of the minute activity during day time (i.e., N=720 being the sequence length, that is, the number of minutes in 12 hours). Before computing the sample entropy of the time series, we applied a Butterworth band-pass filter (independently for each day), whose parameters have been derived with Python library SciPy v.1.1.0. We used a filter of the fourth order and with normalized cut-off frequencies (f_low_ =1/2000 and f_high_ = 1/300). We practically observed that band-pass filtration enables more stable results than that without filtering. As for the DVC^®^ activity (see above), we considered the weekly average RDI throughout the paper.

### Neuromuscular function and body weight assessments

#### Grid hanging test

Mice were tested weekly from the age of 7 weeks. Each mouse was placed in the center of a wire grid (1-cm squares) raised about 50 cm from a bench covered with sawdust bags. After gentle shaking, the grid was rotated upside down and the mouse was left hanging on it. The latency to fall from the grid was recorded over two trials of 60 s each, with an inter-trial interval of approx. 30 min [17]. When the total latency of two trials was under 10 s the test was terminated for that mouse.

#### Grip strength test

Mice of the second cohort were tested on the grip strength meter apparatus (Bioseb, France) every two weeks from the age of 7 weeks until 21 weeks. Each mouse was held gently by the base of its tail over the top of the grid with its torso in a horizontal position, then it was pulled back steadily until it could no longer resist the increasing force and the grip was released [18]. The grip strength meter digitally displays the maximum force applied as the peak tension (in grams) once the grasp is released. For each mouse the grip test consisted of three trials with forelimbs and three trials with all four paws with an inter-trial interval of 60 s. The mean of the three trials was taken as an index of forelimb and all four paws grip strength. Results of four paws measures in this test are in Supporting Information.

#### Body weight, symptoms and humane endpoint assessment

Body weight (BW) was measured weekly from age 7 weeks and after each behavioral test. Data were expressed as percentage variation of weight compared to the first measure taken at 7 weeks of age.

Symptoms like tremors, loss of splay reflex, BW decline, delayed righting reflex were daily monitored at the late stage of the disease and used to determine the humane endpoint (BW loss >20%, complete loss of righting reflex).

### Statistical analysis

Since the same subjects were assessed over time, we performed non-parametric repeated measures analysis on most datasets with Genotype and Sex as between-subject factors and Age (weeks) as within-subject factor (see [14]). We used the rank-based analysis of variance-type statistic (ATS), as implemented in the nparLD R Software package [19]. We performed the statistical tests until age 20 weeks because after that time some subjects were missing. We ran post-hoc analysis where possible, using two-sample T-tests. Since the conventional Bonferroni correction is too conservative for strongly correlated repeated measures (in our case measurements are taken quite frequently with respect to the time scale of development of the disease), we used the D/AP procedure to correct tests for multiple comparisons [20]. This method considers correlation between outcomes and is equivalent to no correction for perfectly correlated measures and to Bonferroni correction for completely independent measures. For reference, we reported tables with different correction methods (no correction, Bonferroni correction, D/AP correction) in the Supporting Information. We performed post-hoc analysis for both DVC^®^ and conventional measures until the age of 20 weeks in which all the cages or mice were still present. In all other cases presented in the paper, not involving repeated measures, we performed two-sample T-test.

We used Python to process and visualize data, which are generally presented as mean ± SEM. We used R to run all statistics (version 3.4.3). We set significance level to α=0.05. For the analysis of DVC^®^ metrics (activity and RDI) the statistical unit is the cage, while for the analysis of physical assessments (BW, grid hanging test, grip strength test) the statistical unit is the individual mouse. Days of cage changing were excluded from the analysis. Moreover, since a male WT mouse died during the experiment at week 16, we excluded its cage from the analysis of DVC^®^ metrics.

## Results

### DVC^®^ monitoring of day and night activity

The DVC^®^ system monitored night and day time activity of mice 24/7 for about 4 months. As expected, mice are more active during the night period [14] and in particular are more active closer to light/dark transitions (Fig 1 and 2).

**Fig 1.**
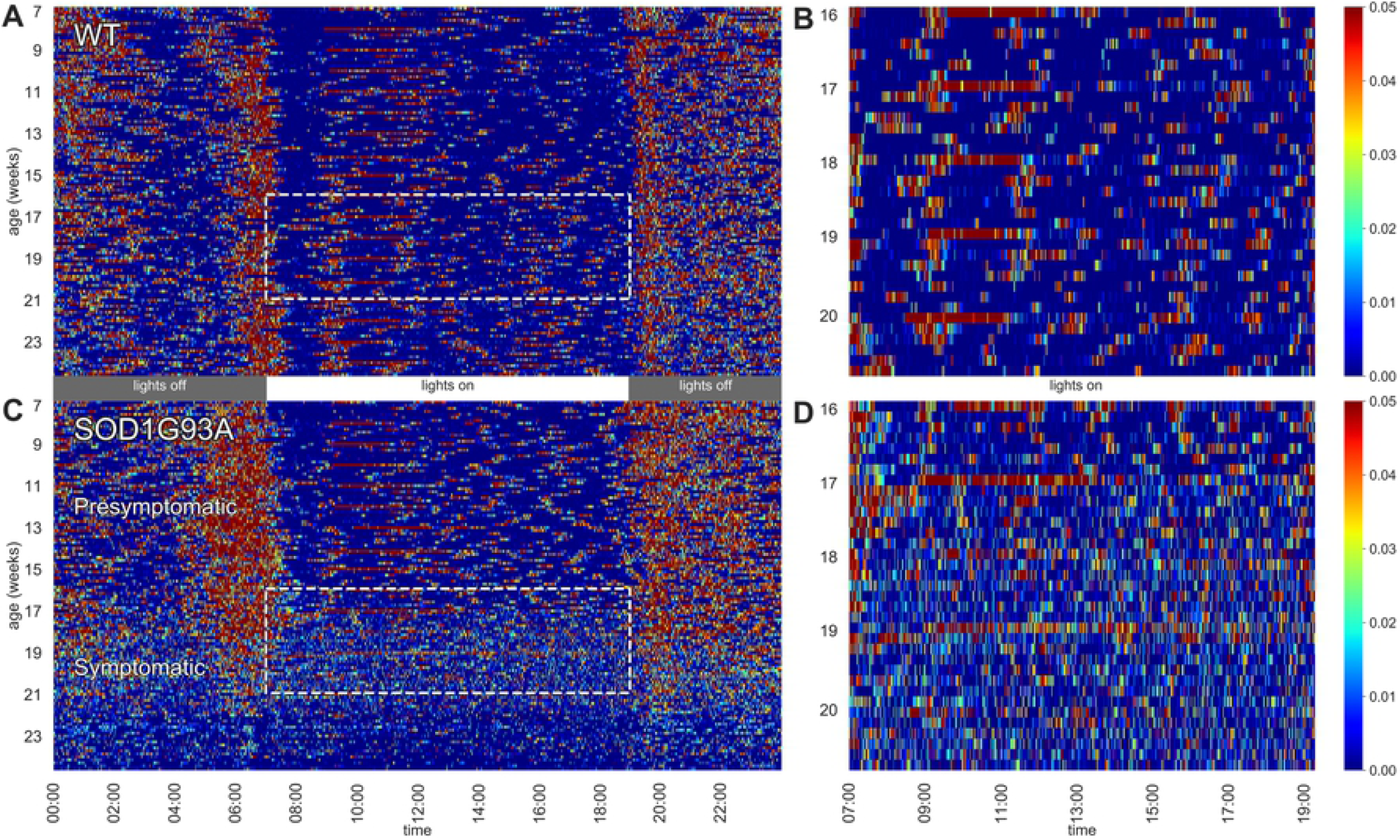
Heat maps of activity. Left panels depict 24 h activity across 7-24 weeks of age in a cage of WT (A) and a cage of SOD1G93A (B) male mice (n=2 per cage). Dotted boxes represent activity during day time between age 16-20 weeks, which are then zoomed in the respective right panels. Weekly cage change related peaks of activity appear clearly as red stripes.

**Fig 2.**
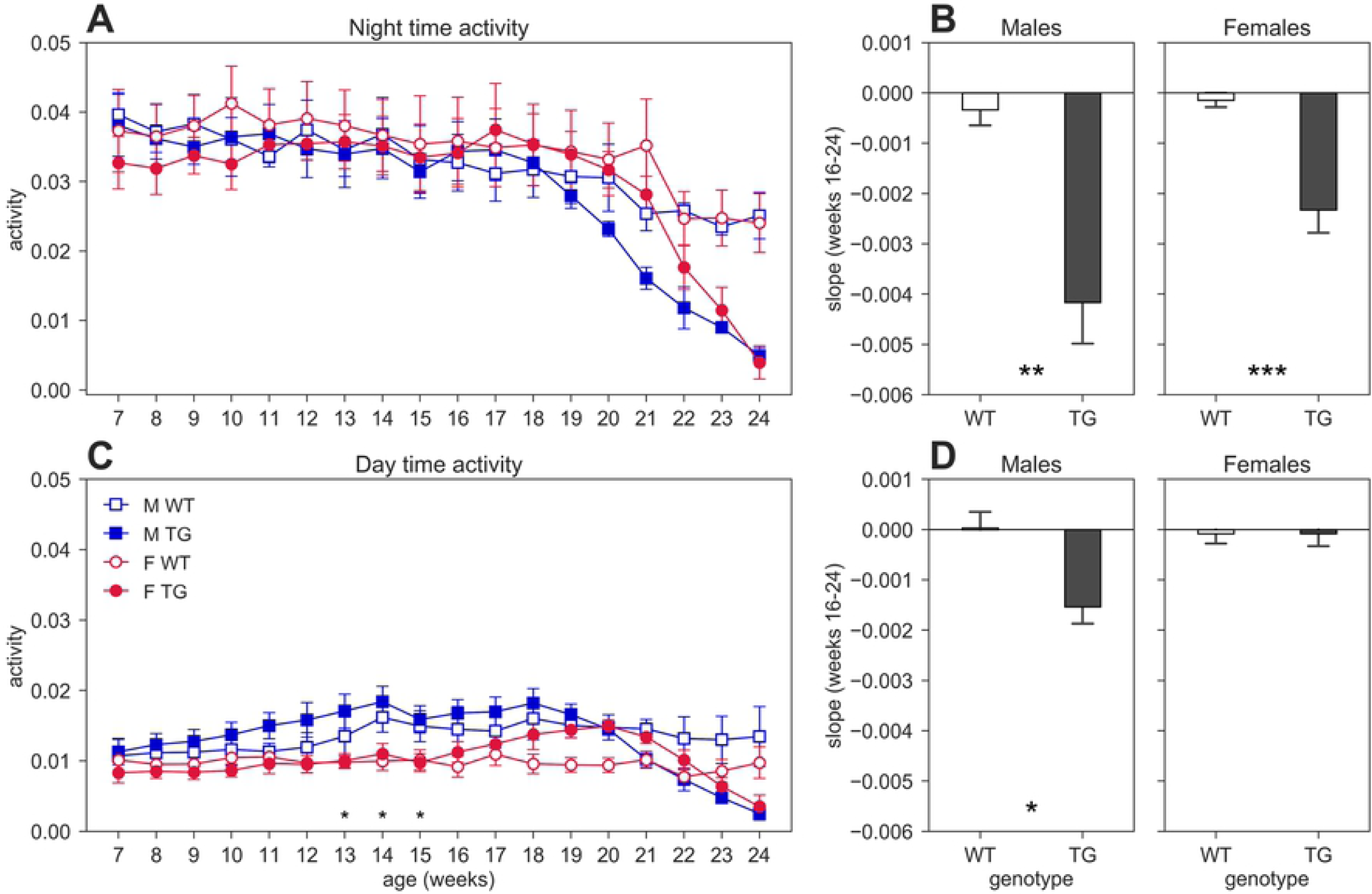
Night and day time activity. Average weekly activity (± SEM) over night (A) and day (C) time across 7-24 weeks of age in male (M, blue squares) and female (F, red circles) cages of WT (open symbols) and SOD1G93A (TG, filled symbols) mice. N of cages per group: M WT=4; M TG=5; F WT=8; F TG=6. We found no significant D/AP post-hoc test when comparing day activity between WT and TG with both males and females. Since we found a significant Genotype X Sex X Age interaction for day time activity (nparLD test, Genotype X Sex X Age interaction: Statistic=3.968, df=5.236, p<0.01) we compared TG males with TG females with D/AP post-hoc procedure in C (*p<0.05, black stars). Panel B shows the slope of the linear regression computed for the weekly night time activity points in weeks of age 16-24, while panel D shows the results for day time activity.

In Fig 1 heat maps of two representative cages of male wild-type (WT) and SOD1G93A (TG) mice (2 mice per cage) show how the activity is distributed across the 24 hours (Fig 1A and C) along the experiment (from age 7 until 24 weeks of age). While the activity of WT mice is very stable across the whole experiment, we observed a clear reduction of nocturnal activity in SOD1G93A cages around 19 weeks of age, corresponding to the symptomatic stage in this mutant strain (tremors, hind limb weakness, overt weight loss). Interestingly, from around 16 weeks of age we observed an atypical pattern of day time activity in the SOD1G93A cages compared to WT littermates (highlighted with the dotted boxes in Fig 1A and C). In Fig 1B and D we zoomed in the day time activity of mice to better appreciate the difference in the activity pattern in a period in which mice usually rest. In Fig 1B the peaks of activity (in red) during routine weekly cage changes are evident.

We measured night and day time cage activity with the DVC^®^ system using the metrics described in [15]. Fig 2 shows the average weekly cage activity during night (Fig 2A) and day (Fig 2C) time. Day activity starts decreasing around age 18 weeks for TG males and age 20 weeks for TG females (nparLD test, Genotype X Age interaction: Statistic=2.595, df=5.236, p<0.05). A similar behavior occurs during night time, where the decreasing trend starts approximately at age 17 weeks. We performed the nparLD test for repeated measures on night activity until week 20 without observing any significant difference across genotype groups.

To further investigate the decreasing trend in TG mice (starting at age 16-17 weeks), we computed the linear regression on the night and day activity between weeks 16 and 24, by considering only weeks with 2 mice in the cage. The slopes of the regression lines are shown in the bar plot of Fig 2B (night) and Fig 2D (day). As expected, the slope for night activity (Fig 2B) for both TG males and females are considerably larger than those of the control mice (two-sample T-test, males: T=3.754, df=7, p<0.01; females: T=4.622, df=12, p<0.001). For day activity, we found a significant difference in the slope only in males (Fig 2D, two-sample T-test, males: T=3.054, df=7, p<0.05; females: T=-0.001, df=12, p>0.05).

### Rest Disturbance Index (RDI)

As introduced in the Methods Section, we observed that TG mice (developing ALS) start showing day time activity patterns that become irregular, with frequent activity bouts, which are not seen in control groups and TG mice at a younger age. The activity anomalies observed in the diurnal pattern of activity in SOD1G93A cages led us to develop RDI as a digital biomarker to quantitatively measure the irregularities of these patterns. Fig 3 clearly shows how this index is remarkably high in both male and female TG mice with a peak around 20 weeks of age, while remaining essentially flat in WT mice (nparLD test, Genotype effect: Statistic=4.648, df=1, p<0.05; Genotype X Age interaction: Statistic=16.637, df=5.463, p<0.001). It appears that this index starts to increase around the age of 11 weeks in male TG mice (Fig 3A) with an ascending curve becoming steeper from weeks 15 to 20 (peak) and then descending when the activity becomes low and some mice were sacrificed. The RDI curve for TG females (Fig 3B) has a sudden increase at week 11 and then remains steady until week 15 (still higher than the control group). Then it increases again with a peak around 20-21 weeks of age and declines similarly to males. Onset of RDI, defined as the age at which TG mice differ significantly from WT (D/AP Post-hoc test), is slightly anticipated in males (age 16 weeks) with respect to females (age 17 weeks) as shown in Fig 3A and B. The average RDI curves for TG males and TG females show similar but shifted patterns (Fig 3C). We thus computed the cross-correlation [21] between the two curves and found that they are maximally similar (i.e., maximum of the cross-correlation) when the curve of TG females is anticipated by 1 week.

**Fig 3.**
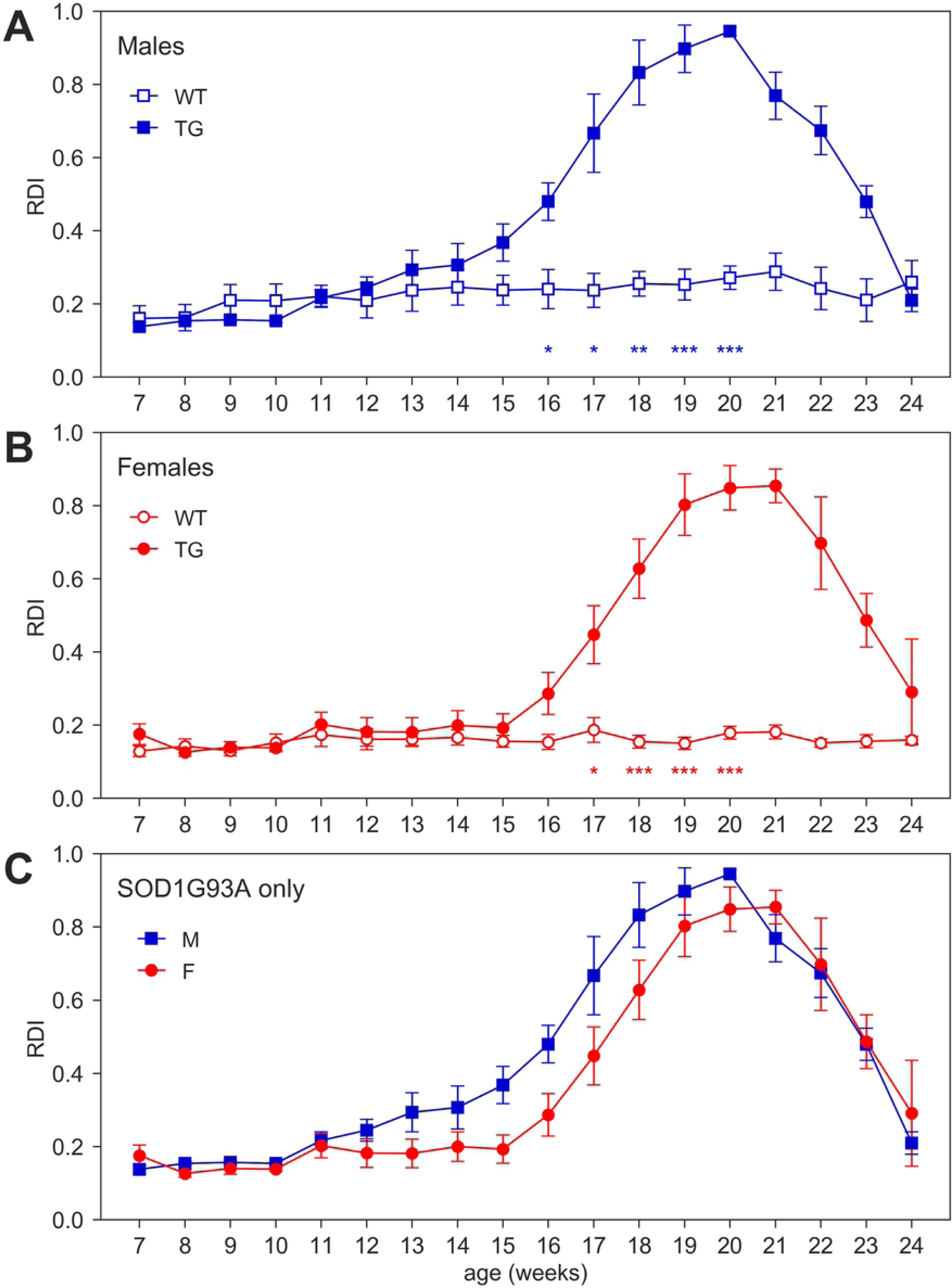
Rest Disturbance Index (RDI). Average RDI curves (± SEM) across 7-24 weeks of age in (A) males, (B) females, (C) TG males and females measured during day time. N of cages per group: M WT=4; M TG=5; F WT=8; F TG=6. In males (A) and females (B) *p<0.05, **p<0.01, ***p<0.001 WT vs. TG, D/AP post-hoc procedure. We found a significant Genotype X Sex X Age interaction (nparLD test, Genotype X Sex X Age interaction: Statistic=2.686, df=5.463, p<0.05) but no significant D/AP post-hoc results when comparing TG males vs TG females.

### Classical ALS model-related measures: neuromuscular function and body weight assessments

Beside the deficit in body weight gain, SOD1G93A mice show the expected decline in neuromuscular function assessed by grid hanging and grip strength tests (Fig 4 and S1 Fig).

**Fig 4.**
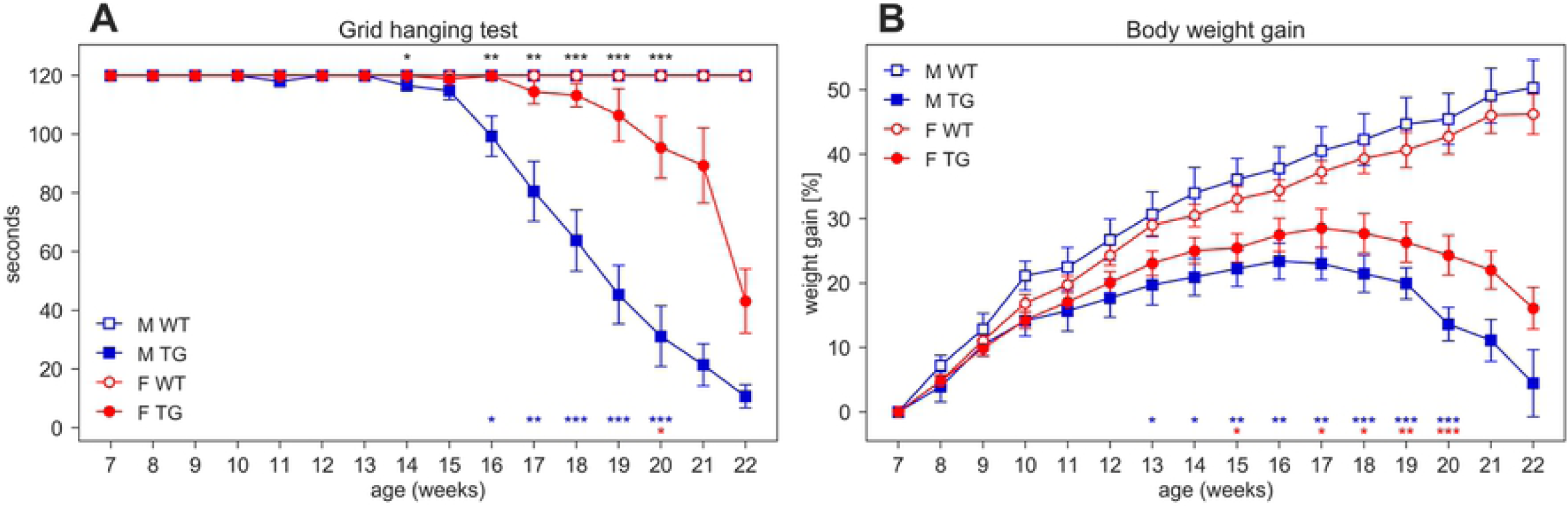
Grid hanging test and body weight gain. (A) Latency to fall per mouse (average time ± SEM) of mice hanging from an inverted grid. (B) Average percentage of body weight gain ± SEM (per mouse) calculated from initial weight at age 7 weeks. Measures were taken weekly (weeks 7-22) in male (M, blue squares) and female (F, red circles) WT (open symbols) and SOD1G93A (TG, filled symbols) mice. N of mice per group: M WT=9; M TG=10; F WT=16; F TG=12. In each sex group (males=blue stars, females=red stars) *p<0.05, **p<0.01, ***p<0.001 WT vs. TG, D/AP post-hoc procedure. Since we found a significant Genotype X Sex X Age interaction in the grid hanging test (nparLD test, Genotype X Sex X Age interaction: Statistic=8.988, df=4.701, p<0.001) we compared TG males with TG females (black stars) with D/AP post-hoc tests in A (*p<0.05, **p<0.01, ***p<0.001).

In the grid hanging test (Fig 4A), the nparLD analysis revealed a significant effect of Genotype, Sex and Age factors and all their interactions (p<0.001). In particular, male TG mice show a significant reduction in grid hanging duration compared to WT starting at 16 weeks of age (D/AP Post hoc tests) while female TG mice show a reduction starting much later. Additionally, male and female TG mice start to clearly differ from each other at age 16 weeks (Fig 4A).

We expressed body weight as percentage of weight gain calculated from initial weight at 7 weeks of age (Fig 4B). While WT mice show a constant weight gain over time, transgenic mice start losing weight around 16-17 weeks of age, moreover TG mice show a slower weight gain since early ages (nparLD test from week 8 to 20, Genotype factor: Statistic=20.293, df=1, p<0.001; Genotype X Age interaction: Statistic=24.533, df=3.758, p<0.001). Male TG mice weigh significantly less than WT starting at week 13 while TG female weigh significantly less than WT starting from week 15 (Fig 4B).

We measured grip strength only in mice of the second cohort (N=29) and, as expected, TG mice showed a decrease in grip strength over time, starting around age 13 weeks (S1 Fig).

### Relation between RDI, activity and classical disease-related measures

We compared DVC^®^-based RDI with average activity (night and day), body weight and grid hanging tests in Fig 5 for TG mice only. Fig 5A compares night activity with RDI for both males and females, showing that RDI starts increasing earlier (age 11 weeks) than the start of the night activity decline (age 16-17 weeks). Similarly, an increase of the RDI does not correspond to a change in the day activity (Fig 5B).

**Fig 5.**
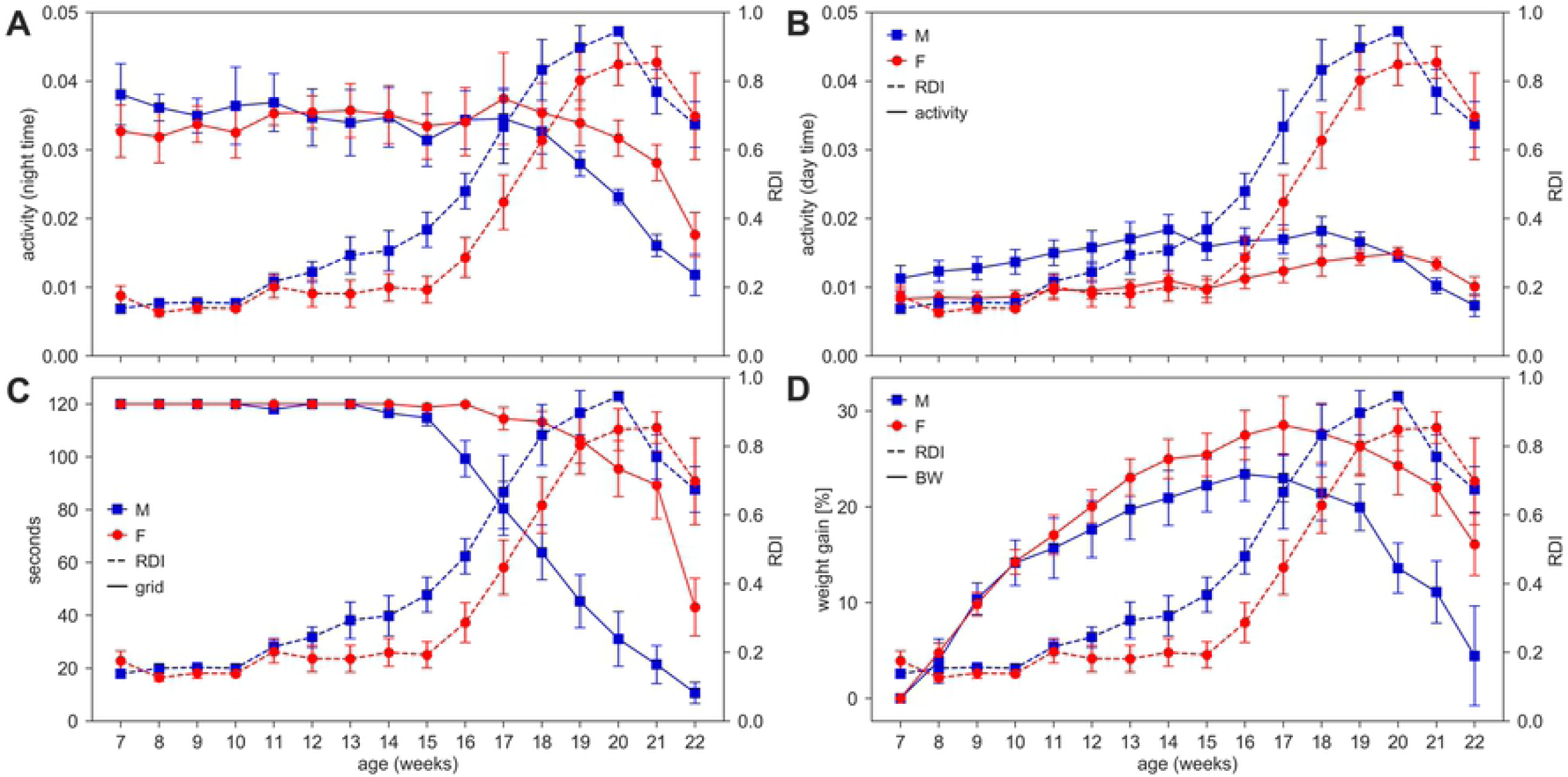
RDI and other parameters in SOD1G93A mice. RDI in cages with SOD1G93A (TG) male and female mice was compared to activity (night and day time, A-B), grid hanging test performance of (C), body weight gain of mice (D). Remember that DVC^®^ metrics are aggregated measures per cage (N: M WT=4; M TG=5; F WT=8; F TG=6), while body weight and grid hanging test are individual measurements (N: M WT=9; M TG=10; F WT=16; F TG=12).

The time course of RDI and grid hanging test latency is shown in Fig 5C, where the decline in grid hanging performance appears at age 14 weeks for males and age 17 weeks for females, later than when RDI starts increasing. Fig 5D shows that body weight gain peaks approximately at age 16 weeks for males and 17 weeks for females, again later than the raise of RDI.

To further investigate the relationship between these measures, we decided to evaluate the Pearson’s correlation between the four metrics of Fig 5 by focusing on the critical interval between weeks 16 to 20 (symptomatic stage). We show the scatter plot between activity during night time and RDI in Fig 6A, where no significant correlation between the two metrics is found, similarly for day time activity in Fig 6B. However, when considering disease-related measurements such as grid hanging and body weight, we found a better correspondence between the increase of RDI and the decline of both grid hanging performance and body weight gain (Fig 6C and D). For grid hanging there is a significant (negative) correlation with the RDI only for males. Moreover, we found significant (negative) correlation between body weight gain and RDI for both sexes, but higher for males.

**Fig 6.**
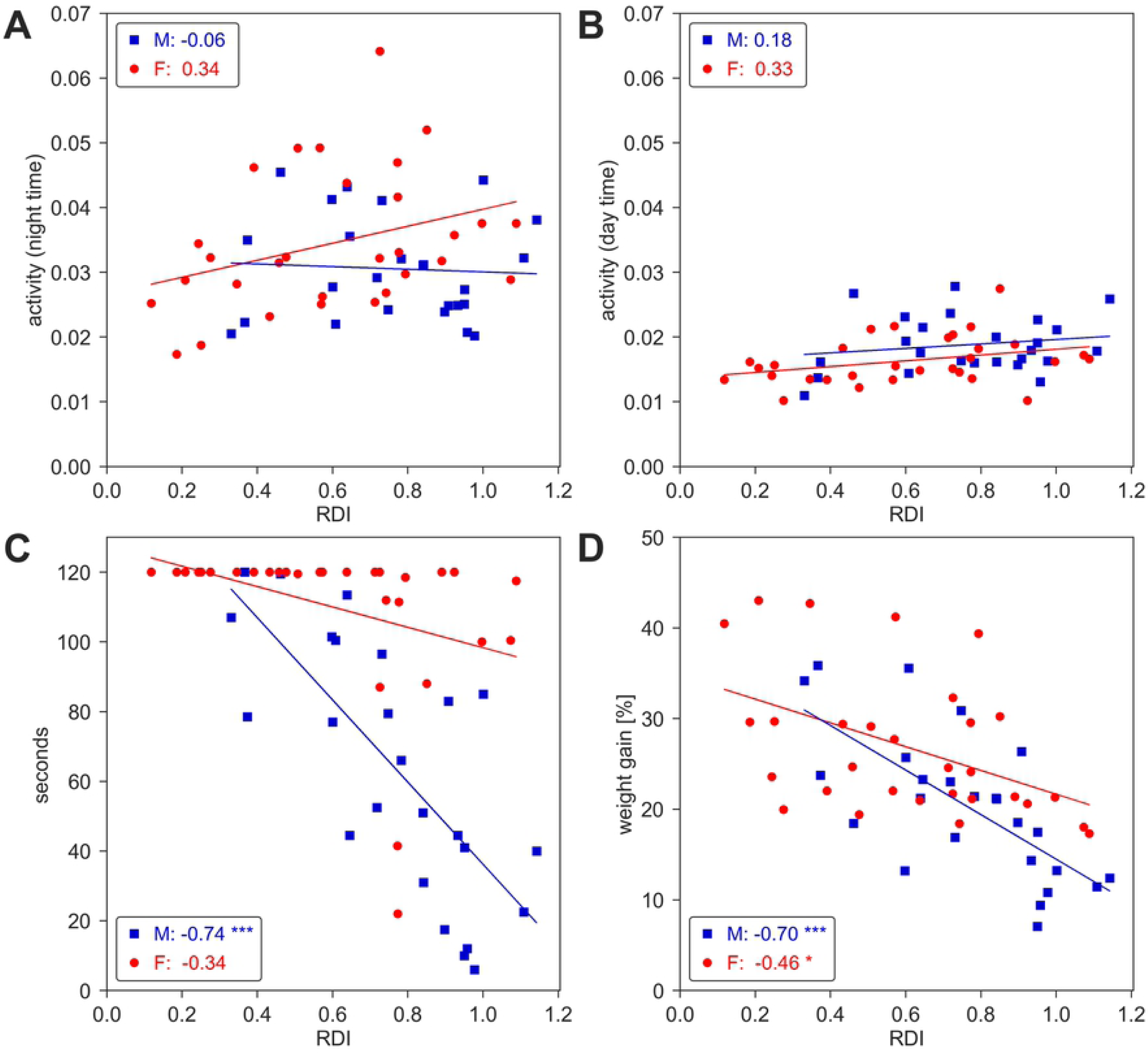
Correlation plots. Correlations between day time RDI and activity (night and day time, A-B), average grid hanging test performance in each cage (C), and average % body weight gain in each cage (D) in male (blue) and female (red) SOD1G93A mice between age 16-20 weeks (M TG = 5 weeks x 5 cages = 25, F TG = 5 weeks x 6 cages = 30 points). Pearson’s correlation coefficient r and significance (*p<0.05, **p<0.01, ***p<0.001) are indicated in each graph for males and females. Regression lines are plotted only for descriptive purposes.

### Effects of husbandry and experimental procedures on activity and RDI

As an additional investigation, we assessed the impact of procedures such as cage change, grid and grip tests, and body weight measurements on activity and RDI metrics. We separated the average activity and RDI measured during weekends (when no procedures are performed) and weekdays in which either cage change or experimental procedures are performed (cage change day or weekdays) (Fig 7).

**Fig 7.**
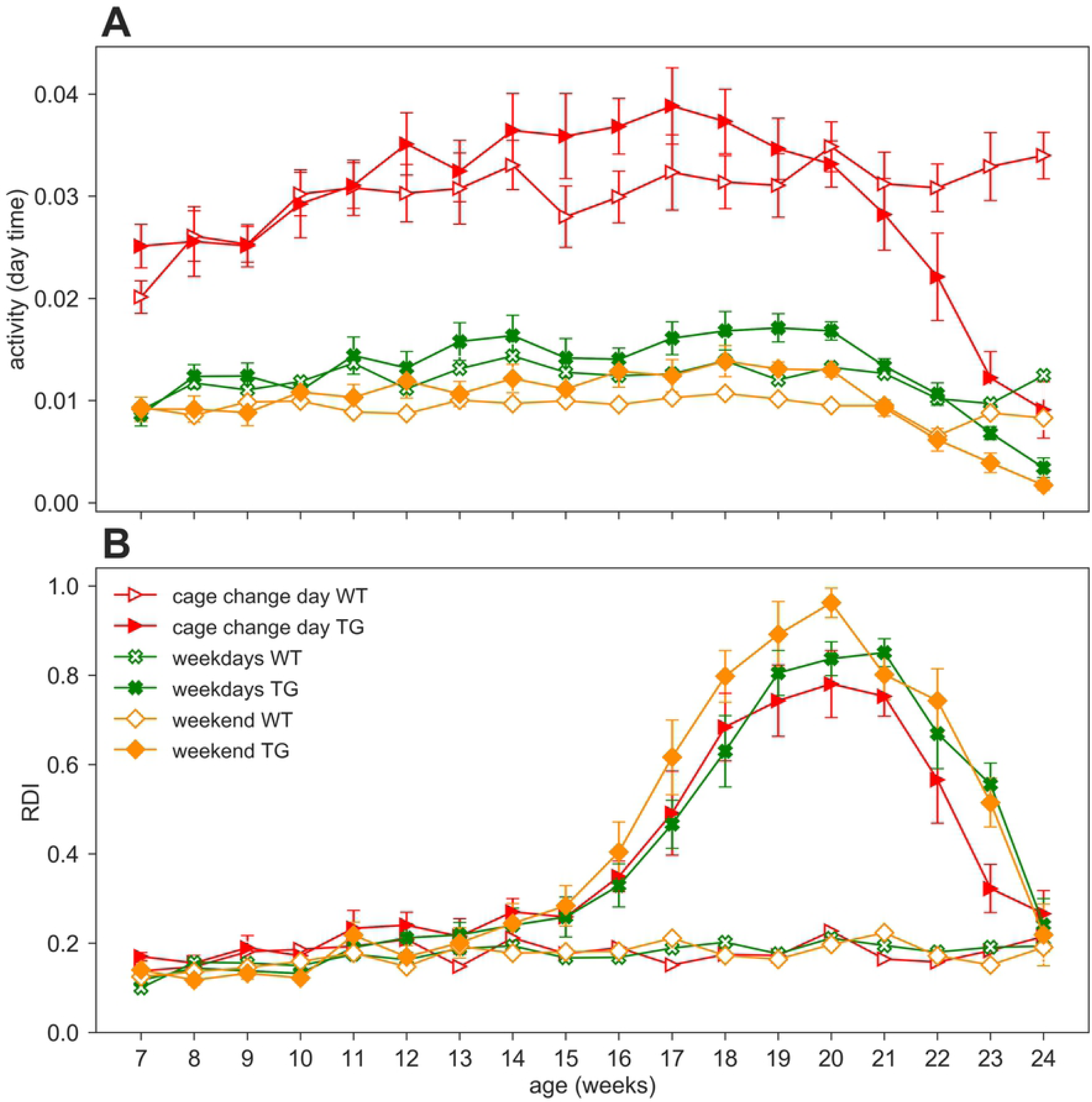
Impact of procedures. Day time activity and RDI curves across weeks 7-24 in WT (open symbols) and TG (filled symbols) (males and females) during cage change days (red triangles), weekdays (green crosses, 2 days in which procedures as grid test or body weight assessment were performed) or weekends (orange diamonds, Saturday and Sunday, without procedures). Data are mean ± SEM. N of cages per group: WT=12; TG=11. No differences were observed in RDI between weekdays and weekends in both genotypes. RDI is not influenced by personnel activity in the room nor by experimental procedures.

We performed the nparLD test with Genotype as between-subject factors and Age (weeks) and Day_type (weekdays or weekend) as within-subject factor. Cage change days as a within-subject factor could not be included in the statistical analysis due to different periodicity in the two cohorts (cage change every two weeks vs. weekly). As expected, day time activity on weekdays is higher than weekends (Fig 7A; nparLD test, Day_type factor: Statistic=117.838, df=1, p<0.001). Day time activity is also strongly impacted by cage change procedures, as already observed in [14], and clearly visible in Fig 7A. For this reason, we remind that days of cage changing were excluded from all the previous analyses.

Interestingly, in RDI does not show a significant difference between weekdays and weekends (nparLD factor, Day_type factor: Statistic=3.527, df=1, p>0.05; Day_type X Genotype interaction: Statistic=0.490, df=1, p>0.05), and similarly the RDI is not substantially affected during cage change days (Fig 7B).

## Discussion

In this paper we presented a novel digital biomarker with the aim to provide a non-invasive method for the early identification of ALS symptoms in male and female SOD1G93A mice. We monitored animals in their home cage via the DVC^®^ system, which is capable of non-intrusively detecting animal locomotor activity 24/7. We considered previously developed metric such as DVC^®^ activity [14,15] and developed a new metric, referred to as Rest Disturbance Index (RDI), which is designed to capture irregularities in the daytime activity pattern. An irregular daytime activity pattern in mice (a nocturnal species), could reflect disturbances in rest/sleep behavior. We also collected measurements about the performance on the grid hanging test, grip strength test and body weight, which are classically used to monitor the progression of the disease [17, 22].

We showed that RDI is a digital biomarker capable of detecting the onset of ALS-related symptoms earlier than pure motor activity and generally earlier than the decline of neuromuscular function as measured in the grid hanging test. The main advantage of RDI is that it is derived from the raw home-cage activity, thus avoiding handling animals compared to other commonly used procedures such as the grid hanging or the grip strength tests. Moreover, RDI is a robust measure that it is not substantially impacted by cage changing or other procedures performed during day time.

One of the main issues in ALS models research is to define unequivocally the onset of the disease and which measure is the most appropriate or reliable. One option is to verify the appearance of significant differences from the control group (WT littermates) in a variety of symptomatic measures (e.g. decline of body weight, neuromuscular dysfunction, coordination deficits and neurological reflexes loss), a second option is to define the age at which those impairments start within the diseased animals groups. The choice of one of the two options or the selection of the test and relevant parameters it is often lab-dependent and there is a lack of consensus on which would be the most appropriate criterion to establish disease onset [10, 23].

In Fig 8, we tried to combine these two approaches comparing the various phenotypic outcomes of SOD1G93A mice to delineate the time course of the disease according to the age at onset of the single parameters. All phenotypes appeared earlier in males than females as already reported in other studies [24]. In the mutant mice the start of change in RDI, grid hanging performance and activity (nocturnal and diurnal), in this order, generally precede the difference with the control group except in the case of body weight gain. In this case the TG mice showed a significantly lower body weight gain compared to WT at an earlier age since they are still gaining weight and have not reached the peak. It is noteworthy that the start increase of RDI appears very early (around 11 weeks of age) compared to the other measures. Additionally, the symptomatic stage that we indicated in the time window 16-20 weeks of age is indeed the interval in which most of the disease related phenotypes emerge. In this critical interval the RDI-related significant difference WT vs TG is in fairly good alignment/correspondence with the decline of night time activity, neuromuscular functions and body weight.

**Fig 8.**
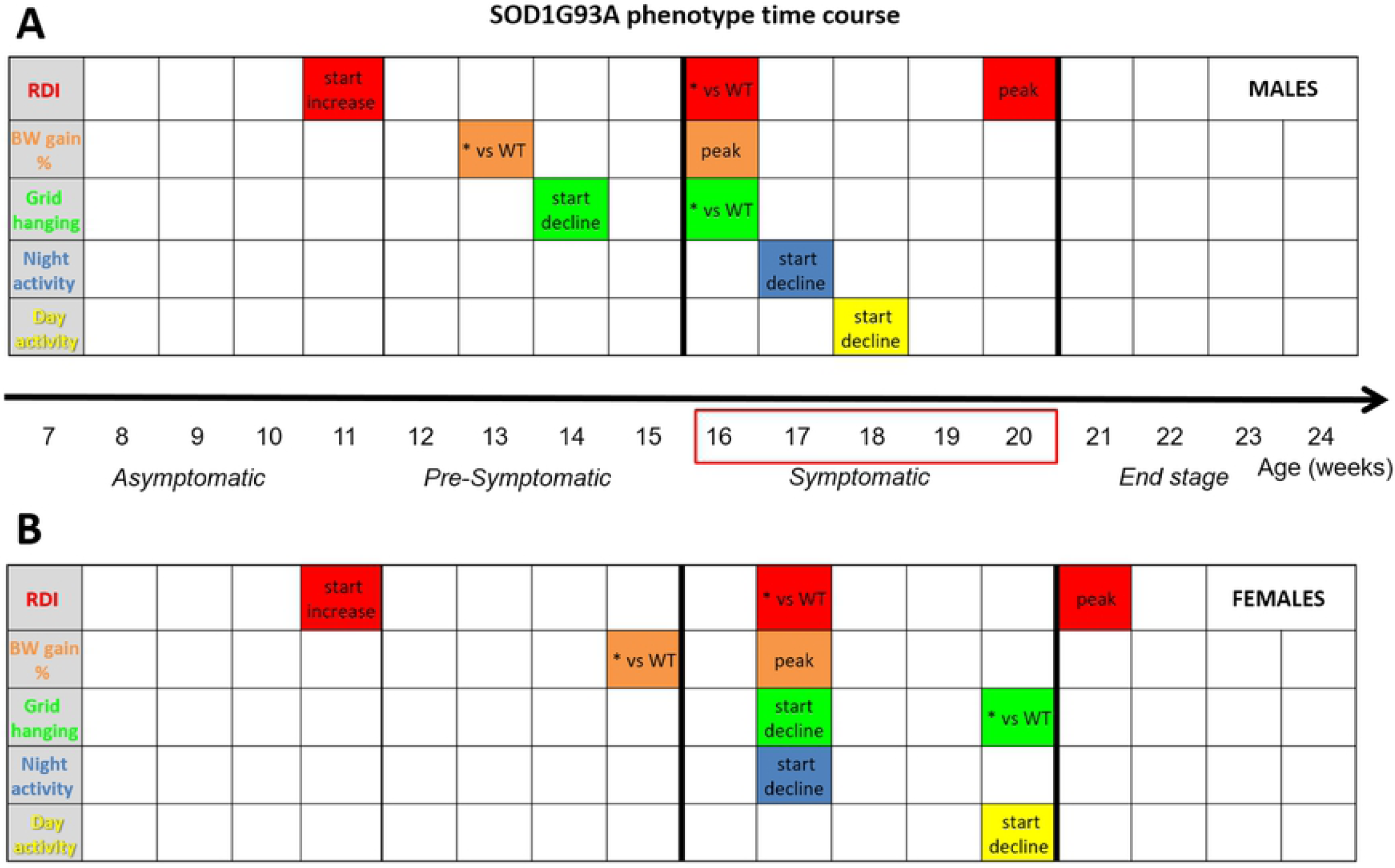
SOD1G93A phenotype time course. Phenotypes listed with their corresponding time points of appearance in SOD1G93A males (A) and females (B) and of statistically significant difference between TG mice and their WT littermates.

In summary, we thus confirm the phenotypes, onsets and the expected sex differences observed in previous studies, by monitoring SOD1G93A mice using both DVC^®^ activity and classical measures. Moreover, we found that the new proposed digital biomarker (RDI) shows a high (negative) correlation with grid hanging performance and body weight gain, especially in males, while no significant correlation with night or day activity in both males and females (Fig 6). Our findings suggest that RDI is a reliable measure of ALS-related sleep fragmentation at the symptomatic stage and also represents an additional marker to detect early signs of the disease starting at a pre-symptomatic stage.

In general, we believe that observing animals in the home cage 24/7 provides clear advantages to experimental data collection, as it allows constant animal monitoring, during both resting and active periods, no animal handling and, crucially, metrics are based on algorithms that are objective and replicable across experiments and sites. This differs from tests performed outside the home cage, which require animal handling during animal resting periods, and often measurements are impacted by operator subjectivity. In addition, we showed that RDI is substantially not influenced by external procedures taking place in the animal room during the day.

The RDI metric, that we suggested being related to rest/sleep disturbances, could be useful as a digital biomarker to early detect disease-related phenotypes also in other neurodegenerative disease models. Circadian rhythm and sleep disruption are in fact recognized as common symptoms that negatively affect the quality of life in many diseases such as Parkinson’s, Alzheimer’s, schizophrenia [4-6, 25] and also ALS [2, 3]. Often these disturbances have been recognized as premorbid signs of the disease and are also typical in the ageing population [26].

In ALS patients, sleep disruption is often caused by respiratory dysfunction and nocturnal muscle cramps and fasciculations accompanied by pain [2, 27].

Very few studies have explored a sleep- or circadian activity-related phenotype in ALS animal models. Sleep fragmentation has been observed in a TDP-43 Drosophila m. model [28]. A study in FUS mutant rats reported sleep and circadian rhythm abnormalities that precede cognitive deficits [29]. In SOD1G93A mice have been observed sleep-related EEG/EMG abnormalities that increase with age [30]. Additionally, always in SOD1G93A mice, melatonin has been shown to increase survival [31] while light-induced disruption of circadian rhythm anticipated disease onset and reduced survival [32]. Our findings, though not measuring sleep directly, agree with a surge of rest fragmentation in both male and female SOD1G93A mice at the symptomatic stage and an indication of early onset of such disturbances.

Overall, we believe that digital biomarkers associated with long-term monitoring of animals affected by neurodegenerative diseases, such ALS, can shed light on behavior and activity patterns that are either unknown or not easily available with conventional, non-continuous animal monitoring. Moreover, the advantages become even more evident when these digital biomarkers can be extracted via technologies capable of non-intrusively monitoring animals 24/7 for several weeks and months. We also envision that digital biomarkers such as RDI, can be successfully applied to other diseases where sleep/rest disturbances appear over time, and that they can be used to preliminary assess the efficacy of treatments in large experiments. In fact, systems such as DVC^®^, can be used to monitor hundreds of cages simultaneously and thus would be very helpful for large-scale mouse phenotyping endeavours as the one undertaken for example by the International Mouse Phenotyping Consortium (IMPC) [33-35].

## Acknowledgments

We thank S. Gozzi and G. D’Erasmo for excellent technical support. We are grateful to the International Mouse Phenotyping Consortium (IMPC) for fruitful sharing of procedures and discussions.

## Supporting Information

**S1 File. Results of grip strength test.** Description of the results of the grip strength test in mice of Cohort II.

**S1 Fig. Grip strength test.** Grip strength (average ± SEM) from 7 to 21 weeks of age. Grip strength performance was assessed every two weeks in male (M, blue squares) and female (F, red circles) WT (open symbols) and SOD1G93A (TG, filled symbols) mice of Cohort II (N=29). N of mice per group: M WT=5; M TG=6; F WT=10; F TG=8. In each sex group *p<0.05, **p<0.01, ***p<0.001 WT vs. TG, D/AP post-hoc procedure.

**S1 Table. Post-hoc analyses with different correction methods (activity, RDI, grid-test, BW).** Summary of p-values obtained from all the performed post-hoc analyses, with different correction methods (no correction, Bonferroni correction, D/AP correction). In red cells, p<0.05.

**S2 Table. Post-hoc analyses with different correction methods (grip strength test).** Summary of p-values obtained from the post-hoc analysis of the grip strength test, with different correction methods (no correction, Bonferroni correction, D/AP correction). In red cells, p<0.05.

